# Decoded EEG Neurofeedback-Guided Cognitive Reappraisal Training for Emotion Regulation

**DOI:** 10.1101/2023.06.10.544438

**Authors:** Linling Li, Xueying Gui, Gan Huang, Li Zhang, Xue Han, Zhen Liang, Zhiguo Zhang

**Affiliations:** School of Biomedical Engineering, Medical School, Shenzhen University, Shenzhen 518060, China; Guangdong Provincial Key Laboratory of Biomedical Measurements and Ultrasound Imaging, Shenzhen 518060, China; Department of Mental Health, Shenzhen Nanshan Center for Chronic Disease Control, Shenzhen 518060, China; School of Computer Science and Technology, Harbin Institute of Technology, Shenzhen 518060, China; Peng Cheng Laboratory, Shenzhen 518060, China; Marshall Laboratory of Biomedical Engineering, Shenzhen University, Shenzhen 518060, China

**Author notes:** These authors contributed equally to this study (Linling Li, Xueying Gui).

**Keywords:** decoded neurofeedback, electroencephalogram (EEG), emotion classification, cognitive reappraisal, emotion regulation

## Abstract

Neurofeedback is an innovative self-training that adjusts and enhances brain function. One of the more studied application scenarios with neurofeedback training is emotion regulation. Compared with regional brain activity-informed neurofeedback techniques, neurofeedback protocols using the decoded brain states as feedback signals could make the setting of regulation targets more specific and objective. In the present study, we constructed an EEG neurofeedback-guided cognitive reappraisal training protocol for emotion regulation. Forty-two healthy participants (20 females; 22.4±2.2 years old) were recruited and were randomly assigned to either the neurofeedback group or the control group. During the training process, we calculate the real-time self-regulation performance on the evoked emotion based on the decoded emotional states and fed it back to the subjects as the feedback signal. According to our results, real-time feedback of the regulation effect helps subjects improve emotion regulation performance for emotional stimuli with low positive valence. Further analysis of selected EEG features for emotion classification revealed the neural correlates with neurofeedback training. This newly proposed neurofeedback training protocol is a promising treatment for emotion-related mental disorders, with the potential to be a low-cost and high-portability brain-based, non-invasive, neural modulation technique.

## 1 Introduction

Neurofeedback is an innovative tool in which neural activity is measured, and representation of this activity is presented to the participant in real-time for learning self-regulation of the putative neural substrates that underlie a specific behavior of pathology (Sitaram *et al*., 2017). Neurofeedback training approaches are usually based on real-time measures of brain activity using neuroimaging techniques, with the majority of applications relying on electroencephalography (EEG) and some employing functional magnetic resonance imaging (fMRI). One of the more studied application scenarios with neurofeedback training is emotion regulation. Generally, emotion regulation is a crucial skill associated with well-being and mental health (Gross, 2002). One potential mechanism in the development and maintenance of psychopathology, such as depression, is difficulties with emotion regulation (Gross and Jazaieri, 2014; Donofry *et al*., 2016). Therefore, improving emotion regulation skills is a primary aim of psychotherapeutic interventions and neurofeedback training is a promising brain rehabilitation technique (Sloan *et al*., 2017).

### 1.1 Regional Brain Activity Informed Neurofeedback

Existing studies have applied neurofeedback by measuring regional fMRI activation or EEG oscillations for emotion regulation training. For example, healthy people can regulate their brain activity in the presence of real-time fMRI neurofeedback from various brain regions related to emotion regulation, including the amygdala, anterior insula, and anterior cingulate cortex (Linhartová et al., 2019). Compared with fMRI neurofeedback, EEG neurofeedback has the advantage of being more widely available, cost-effective, and including ambulatory settings. Existing EEG studies for emotion regulation used single-frequency band EEG activity in specific brain regions for neurofeedback signal calculation, such as alpha symmetry (Quaedflieg *et al*., 2016; Mennella *et al*., 2017; Li *et al*., 2023), and high-beta down-training (Paquette et al., 2009; Cheon et al., 2016; Wang et al., 2019). However, the generation, perception, and regulation of emotions depend on the synergistic work of multiple brain regions or brain networks (Morawetz et al., 2020). These regional brain activity-informed neurofeedback techniques for emotion regulation might have limited capability for induced learning effects during training.

### 1.2 Decoded Brain State Informed Neurofeedback

With the help of machine learning techniques, emotional states can be decoded from human brain activity, and the decoded brain states can be used as the neurofeedback signal to make the setting of regulation targets more specific and objective (Shibata et al., 2019). Most of the previous decoded neurofeedback studies were performed using fMRI (Watanabe et al., 2017). Based on the same consideration, Bu et al. used decoded brain state associated with smoking cue reactivity as the neurofeedback signal to construct a novel EEG neurofeedback protocol for nicotine addiction, which produced short- and long-term effects on cigarette craving and smoking behavior (Bu et al., 2019). Additionally, Huang et al. developed a neurofeedback training method during which the subjects’ current decoded emotional states are provided to them through visual feedback in order to help them regulate their emotion toward a specific emotional state (Huang et al., 2021). These two studies verified the feasibility of the construction of decoded neurofeedback protocol using a real-time collection of EEG signals and brain state decoding.

### 1.3 Cognitive Reappraisal for Emotion Regulation

Cognitive reappraisal is one of the most effective emotion regulation strategies and is the effortful modification of a situation’s meaning to change its emotional impact (Gross, 2002; Gross, 2015). Empirical evidence consistently suggests that cognitive reappraisal is beneficial for psychological health in clinical settings (Forkmann et al., 2014). One particular challenge which may impede the acquisition of cognitive reappraisal strategies through behavioral therapy is that the subject may lack meta-cognitive awareness of the impact of psychotherapeutic strategies. To address this issue, a previous study used real-time fMRI collection and provided neurofeedback during cognitive reappraisal training, and their results suggested that the real-time feedback of regulation performance could help subjects learn to voluntarily self-regulate their brain signal in the brain circuitry of emotion regulation and facilitate the application of regulatory strategies real-world situations in daily life (Zweerings et al., 2020). Therefore, developing cognitive reappraisal training protocol based on EEG neurofeedback may offer more convenient neuroscience-based interventions for self-regulation training.

### 1.4 Work in the Current Study

In the present study, we, therefore, constructed an EEG neurofeedback-guided cognitive reappraisal training protocol and evaluated the training effects in healthy subjects. More specifically, each participant finished one neurofeedback training session, which consisted of a calibration run and two emotional regulation runs for emotional stimuli with different valence levels (low positive and high negative). First, a personalized two-classes emotion classifier was trained using data collected during the calibration run. Then, during regulation runs, participants were trained to increase the positive affect or reduce the negative affect elicited by affective picture stimuli using cognitive reappraisal strategies. Besides, feedback of self-regulation performance was provided in real-time to help participant adjust their strategies. This study employed a single-blind design and the enrolled 42 subjects were randomly assigned to either the neurofeedback group (n=21) or the control group (n=21). We aimed to determine the neurofeedback training effect and make comparisons between experimental groups, as well as investigate the evoked brain activity changes underlying brain mechanisms of emotion regulation.

## 2 Material and Methods

### 2.1 Participants

Forty-two healthy participants (20 females; 22.4±2.2 years old) were recruited through an online advertisement at Shenzhen University. The participants were all right-handed according to the Edinburgh Handedness Inventory and had normal or corrected-to-normal vision. Participants were excluded for any history of psychiatric conditions (e.g., depression), neurological disorders (e.g., stroke), or psychotropic substance use. None of the subjects had prior experience with neurofeedback-training experiments. The study was approved by the Medical Ethics Committee of Shenzhen University (approval number: 2019017). Written informed consent was obtained from all the participants before the experiment.

### 2.2 Emotional Stimuli

This study used pictures for emotion elicitation. Two hundred and forty color pictures were selected from the Chinese Affective Picture System (CAPS) datasets (120 positive pictures with high valence, and 120 negative pictures with low valence) that represented a range of content as assessed by the reported norms for valence and arousal (Lu et al., 2005). Compared with negative pictures, normative ratings for positive pictures were more present (positive: 6.62±0.60; negative: 3.10±0.94; p < 0.001) and they were matched regarding arousal ratings (positive: 5.92±0.65; negative: 5.28±0.44; p < 0.001).

### 2.3 Neurofeedback Training

The experiment included one training session, which consisted of a calibration run and two emotional regulation runs. During the experiment, the subjects were seated in front of a computer monitor and were instructed to stay as still as possible to minimize body movement artifacts. Participants were randomly assigned to either the neurofeedback group (n=21) or the control group (n=21) in this single-blind study. During the emotional regulation run, the feedback signal was only displayed for the neurofeedback group.

#### 2.3.1 Calibration Run

In the calibration run, the subjects were exposed to two types of emotional stimuli (120 pictures of each category), and the task was divided into three runs. Each trial began with an image presentation for 3000 ms, during which participants were required to watch passively. They were then asked to report their level of emotional valence on a 10-point scale (a high score indicated a high level of positivity) within 1500 ms using the keyboard. The interstimulus interval (ISI) of 500–1000 ms (pseudorandomized within each run). When the calibration run was finished for each subject, a two-class linear support vector machine (SVM) classifier was trained using the recorded EEG data and subjective ratings, which were used in the following emotional regulation run.

#### 2.3.2 Regulation Run

After the calibration run, each subject performed two neurofeedback training runs of emotional regulation. Based on the subjective ratings of emotional valence recorded in the calibration run, 30 pictures with low positive valence (6.30±0.39; LP regulation run) and 30 pictures with high negative valence (0.98±0.86; HN regulation run) were selected for each subject. Then two regulation runs were completed for each set of selected pictures in order to compare the neurofeedback training performance between eliciting greater positive emotions and lesser negative emotions. The order of the two regulation runs was counterbalanced.

Each regulation run consisted of six pairs of blocks, one passive view block and one following self-regulation block, each containing five trials. Each pair of blocks consisted of the same five pictures but presented in a different order. Before each block started, a cue lasting 2s was displayed on the screen, indicating “passive view” or “self-regulation”. In the passive view block, each picture was presented for 3000 ms with an ISI of 2000–2500 ms. In the self-regulation block, after the picture presentation and an ISI of 4000-5000 ms, the feedback signal showed for 2000 ms reflecting the extent to which the brain activity matched the pattern for a positive emotional state. Here we implemented intermittent feedback to reduce dual-task interference, and a recent study indicated that intermittent feedback may be superior for shaping brain activity compared to continuous feedback (Hellrung et al., 2018). The details of feedback signal calculation will be introduced in the data analysis section. Another ISI of 500-1000 ms was presented following the neurofeedback signal. For emotional self-regulation, subjects were instructed to use cognitive reappraisal strategies to regulate elicited emotions with two types of goals. All participants received instructions on cognitive reappraisal strategies before the experiment of regulation runs (Goldin et al., 2008). The participant’s goal during the LP regulation run was to upregulate the positive emotional response through either engaging with the situation, reinterpreting the situation in a more positive way, or perceiving the situation as being more real. The participant’s goal during the HP regulation run was to downregulate the negative emotional response through either reinterpreting the situation in a less negative way, observing detachedly, becoming psychologically distant, or looking at it in a more objective way.

The affective state was measured with the Positive and Negative Affect Scale (PANAS) (Schmukle et al., 2002) before and after the regulation runs. A neurofeedback training effect questionnaire was used to measure participants’ subjective feelings and strategies. Three questions were included in the questionnaire: i) Please rate the level of fatigue after the completion of the experiment (1-9 points); ii) Please evaluate the degree of concentration during training (scale 1 to 5); iii) Please evaluate whether the strategy you use is effective (scale 1 to 5); iv) Please briefly describe the cognitive reappraisal strategies used during neurofeedback training.

### 2.4 EEG Data Acquisition

EEG signals were recorded using a BrainAmp amplifier (Brain Products GmbH, Germany) with 64 Ag/AgCl electrodes placed according to the extended international 10-20 system. Channels were recorded with a reference at FCz and a common ground at FPz. EEG signals were recorded in the BrainVision recorder software (Brain Products GmbH, Germany). The sampling rate was 1000 Hz for EEG data acquisition during calibration runs and resting-state runs, and 500 Hz for online acquisition during regulation runs. The impedance of all electrodes was kept below 20 kΩ. EEG signal was collected during neurofeedback training (one calibration run and two regulation runs) and resting-state (2 min eyes-open and 2 min eyes-closed) before and after training each regulation run.

### 2.5 EEG Data Analysis

#### 2.5.1 Preprocessing

The EEG raw data preprocessing was conducted by the EEGLAB toolbox for MATLAB (https://sccn.ucsd.edu/eeglab). The preprocessing steps included: (1) removal of the IO channel; (2) 1-45Hz band-pass filtering and 49-51Hz notch filtering; (3) downsampling to 500 if the raw data was recorded at a 1000 Hz sampling rate; (4) segmentation of epochs (500 ms before and 3000 ms after stimulus onset); (5) baseline correction using the pre-stimulus interval; (6) independent component analysis (ICA) (Bell and Sejnowski, 1995) to remove artifacts embedded in the data (e.g., muscle, eye blinks) based on IClabel (EEGLab plugin) (Pion-Tonachini et al., 2019); (7) common average reference.

#### 2.5.2 Feature Extraction

We extracted the time domain and frequency domain features which have been widely used for classifying emotions in existing studies from the EEG signal of each epoch from each channel (Rahman et al., 2021). Hjorth parameters (activity, mobility, and complexity) were calculated from the non-linear analysis in the time domain (Hjorth, 1970; Li et al., 2018; Joshi and Ghongade, 2021). We also extracted the amplitudes of N1 (130-180ms), P1 (80-130ms), N2 (180-280ms), P2 (180-280ms), P300 (280-400ms), and LPP (400-500ms) of ERPs in the time domain as features that are related to emotions (Ding et al., 2017; Singh and Singh, 2017). Power spectral density (PSD) was estimated using Welch’s algorithm (1024-point FFT, 50% Hanning window), and power features were extracted in five frequency bands [delta (1-4 Hz), theta (4 -8 Hz), alpha (8-13 Hz), beta (13-30 Hz), and gamma (30-45 Hz)] (Rahi and Mehra, 2014; Wang et al., 2014; Gao et al., 2020). The di□erential entropy (DE) was calculated for each frequency band and it was equivalent to the logarithmic PSD (Shi et al., 2013; Zheng et al., 2019; Patel et al., 2021). The discrete wavelet transform (DWT) was performed for feature extraction in the transform domain (Wang et al., 2014; Mohammadi et al., 2017). DWT analyzes signals in different frequency bands at different resolutions by decomposing the signal into coarse approximations and detailed information. Here we used Daubechies 4th order because of good time-frequency properties and four decomposition levels wavelet energy features were extracted (D4, gamma; D5, beta; D6, alpha; D7, theta). After concatenating EEG features independently obtained from 63 EEG channels to form a single feature vector, the final dimensionality is 1701 for each data segment.

#### 2.5.3 SVM Model Training

The data samples extracted from each calibration run were used to train the SVM model for emotion classification, which was subsequently used for online prediction in the regulation runs. We perform feature selection based on the Eigenvector Centrality Feature Selection (ECFS) algorithm (Roffo and Melzi, 2016), which generates a graph of features with features as nodes and evaluates the importance of each node through an indicator of centrality, i.e., eigenvector centrality.

We utilized the linear SVM algorithm and 10-fold cross-validation to compare the classification performance of the feature subsets with different feature dimensions (N = 6, 11, 16, …, 1701), and the top K features with the best classification accuracy were selected as the optimal feature subset. With each iteration, each feature was normalized to the subjects in the training sample by subtracting the minimum and dividing by the difference of the maximum minus the minimum, and the same maximum and minimum values were used to scale the test data. Subsequently, based on the determined K value, hyperparameter optimization was performed for the ECFS algorithm, a is taken to 1 with 0 as the initial value and 0.01 as the step size, and the classifier construction under ten-fold cross-validation is carried out, and finally, the a value with the highest average decoding rate is obtained. Finally, based on the determined K value and a value, the final prediction accuracy was evaluated using 10-fold cross-validation, and one trained classifier model for each subject was obtained using all the data samples from the calibration run.

We also explored the selected EEG features in with-subject emotion classification from a group-level perspective and multiple perspectives, including feature types and channel locations. Specifically, for each feature at each channel location, we calculated the percentage of subjects who have selected the current feature and it could indicate the degree of robustness of the feature at group-level. Among the 27 feature types, the features that had a greater percentage value than average were selected as the group-level discriminating features for emotion classification and were used for further analysis. Seven Regions of Interest (ROIs) were determined based on the 63 channels of the acquired EEG signals (Wang et al., 2014; Zheng and Lu, 2015; Zhuang et al., 2018; Zheng et al., 2019): the left anterior frontal ROI 1 (including Fp1, AF7, and AF3), the right anterior frontal ROI 2 (including Fp2, AF8, and AF4), left temporal ROI 3 (including FT9, FT7, T7, TP9, and TP7), right temporal ROI 4 (including FT10, FT8, T8, TP10, and TP8), and frontal Central ROI 7 (including FC1, FC2, CZ, and Fz), left Parietal ROI 5 (including P5, P7, and PO7), and right parietal ROI 6(including P6, P8, and PO8) (as shown in Figure 4C). For the group-level discriminating features, each category of selected EEG features in each ROI region was averaged, and summarized feature values of this ROI region were calculated.

**Figure 1.**
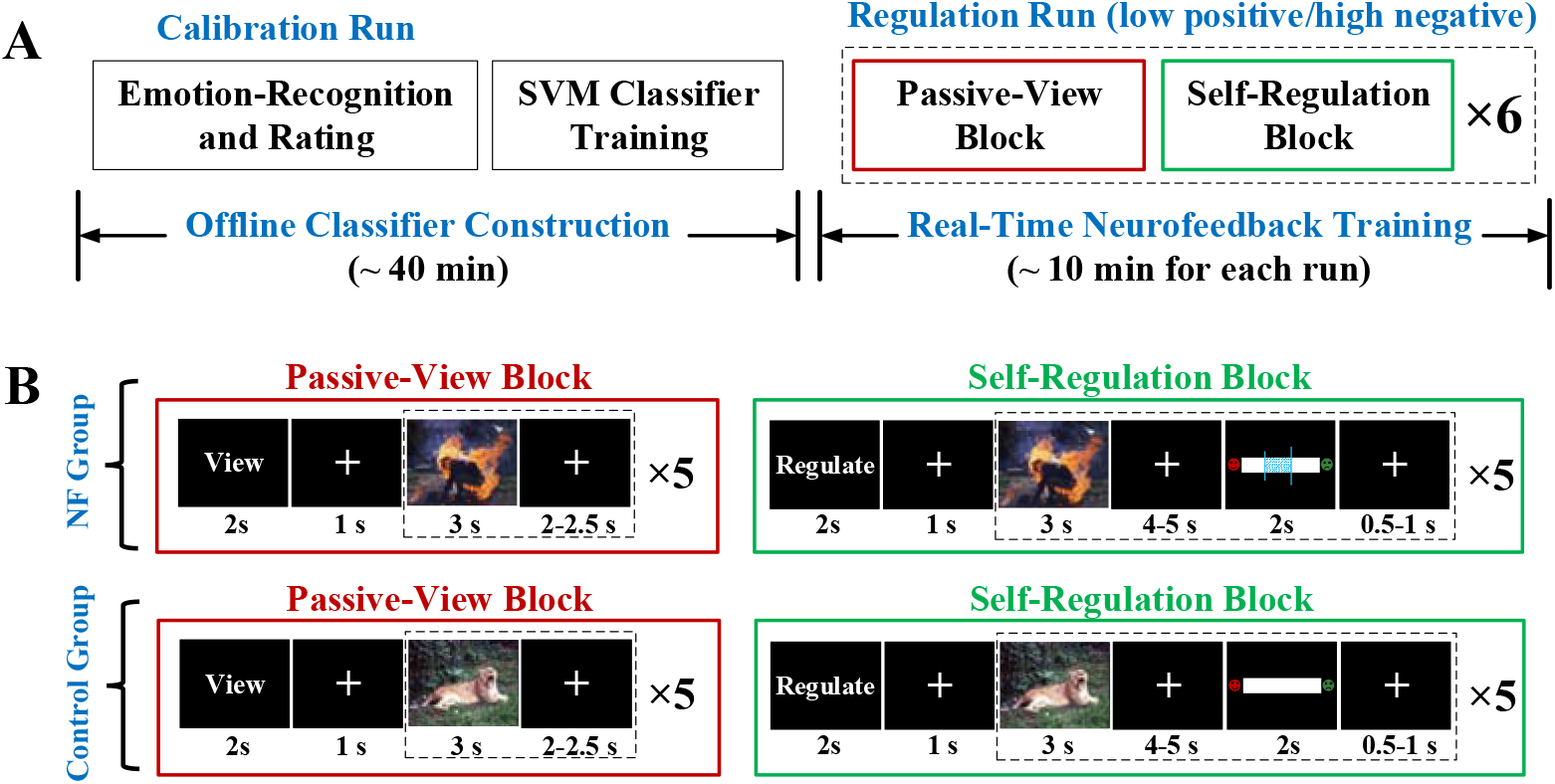
Experimental procedure and neurofeedback paradigm. A. The experimental procedure consisted of two parts: one calibration run for offline emotion classifier construction and two neurofeedback training of emotion self-regulation runs (picture stimuli with different levels of emotional valence). Each regulation run consisted of 6 pairs of passive-view blocks and self-regulation blocks (same pictures but presented in a different order). B. Paradigms of neurofeedback training. Each block consisted of 5 trials. On ‘view’ trials, participants were asked to respond naturally to each picture; on ‘regulation’ trials, participants had to reappraise the pictures in order to increase the elicited positive affect or reduce the elicited negative affect. The neurofeedback signal, calculated as the changes of decoded emotion state was displayed after each ‘regulation’ trial only for the neurofeedback (NF) group.

**Figure 2.**
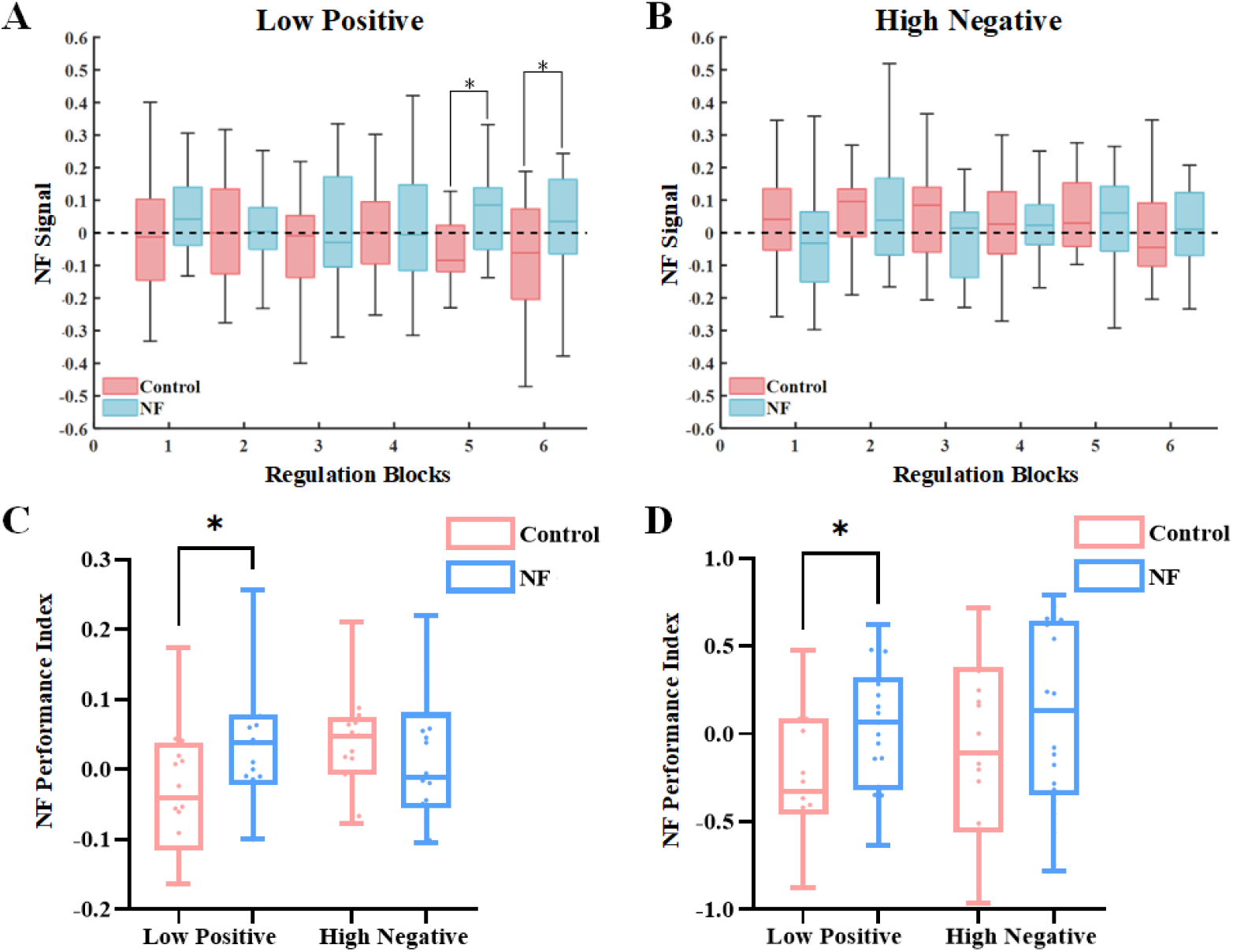
Neurofeedback learning performance of emotion regulation. *p < 0.05; error bar (SD); NF: neurofeedback.

#### 2.5.4 Online Prediction and Feedback Calculation

During the regulation runs, the online EEG data preprocessing and feature extraction were the same as the previous offline data analysis for the calibration run. For the trials with the same stimulus picture in the passive view block and self-regulation block, EEG features were extracted (top K features with the best classification accuracy) and fed into the trained classifier from the calibration run separately. The classifier estimated the probabilistic scores (from 0 to 1) in real-time, when the current activity patterns are more similar to the positive emotional state, the score increased. Then the calculated probability scores for the same stimulus picture in the passive view block (short line) and self-regulation block (long line) were displayed in the form of horizontal bars as the feedback signal. Thus, by comparing the position of the probability scores, the subject was informed of the self-regulation performance on the evoked emotion by the current stimulus picture. The subjects were instructed to increase the change of probability score in the positive direction by adjusting the cognitive reappraisal strategies.

#### 2.5.5 Evaluation of Neurofeedback Training

The index of neurofeedback learning performance was calculated as the mean of the neurofeedback signals across all trials in the self-regulation block and it could reflect the overall amount of change in the emotional state due to self-regulation.

### 2.6 Statistical Analysis

The chi-square test was used for gender and independent two-sample t-tests were used for age and psychological ratings of behavioral scales which were only rated once. The mixed effects ANOVA (between-subject factor: NF group vs. Control group; within-subject factor: before vs. after training) was computed on the PANAS ratings separated for the positive and the negative subscale. The mixed two-way ANOVA was also performed to compare the subjective ratings of the emotional valence of picture stimuli used in the regulation runs and the calculated indices of neurofeedback training performance. Post-hoc tests with Bonferroni’s correction were used to follow-up significant main effects. A two-sample t-test was performed to compare the prediction accuracy of the emotion classification model constructed using EEG data of calibration run between NF and Control groups.

Average values of the selected EEG features (averaged within seven ROIs) were first calculated by averaging all the view trials and regulation trials separately for each training run. Paired sample t-tests were then applied to make a comparison of EEG features between the view block and the regulation block.

Correlation analyses were performed between neurofeedback performance and brain activity change. Firstly, paired t-tests were performed to compare the EEG features of the two groups’ seven ROIs before and after emotion regulation, and then the amount of change in the features before and after emotion regulation was calculated separately for the two groups based on the significantly different EEG features. Bonferroni correction was applied to counteract the multiple comparisons among ROIs (P<0.05/7). Pearson correlation analysis was performed between them and the neurofeedback training performance.

## 3 Results

### 3.1 Demographic and Behavioral Ratings

The NF group and control group did not differ with respect to gender, age, stage of depression (PHQ-9), anxiety (GAD-7), and trait emotion regulation (ERQ) (all p>0.05). According to the mixed ANOVA using the pre- and post-test scores of PANAS, there was no significant main effect of group (positive subscale, p = 0.466; negative subscale, p = 0.750), nor a significant interaction effect (positive subscale, p = 0.330; negative subscale, p = 0.197), but a significant effect of training (positive subscale, p = 0.002; negative subscale, p <0.001), indicating both increased positive and negative effect after training for both groups. No group difference was observed for the self-reported subjective feeling of fatigue, how concentrated, and neurofeedback learning success (all p>0.05). According to the mixed ANOVA using emotional valence of picture stimuli used in the regulation runs, there was no significant interaction (p = 0.855), no significant effect of group (p = 0.468), and only a significant effect of regulation type (p <0.001).

### 3.2 Neurofeedback Performance

We assessed the neurofeedback learning performance for evoked changes of emotional states decoded from brain signals. The index of learning performance was calculated for each subject, and it reflected the overall amount of change in the emotional state. A mixed two-way ANOVA on the performance index L1 identified a significant interaction effect between experimental grouping and regulation type (p=0.005), and no significant main effect of group (p=0.361) and regulation type (p=0.091). As further indicated by the results of post hoc tests with Bonferroni adjusted significance level, there was no significant difference in the L1 of the regulation effect index between the feedback and control groups (p=0.419) under the High Negative regulation type. But there was a significant difference in the L1 of the regulation effect index between the two groups of participants (p=0.031) under the Low Positive regulation type.

### 3.3 Emotion Decoding Accuracy and Selected Features

The accuracies of the constructed emotion decoders for online prediction in the regulation run are shown in Figure 3. The classifier was obtained using the entire sample data collected in the emotion recognition and rating task. The mean accuracy for the NF group was 68% (range: 62%-74%), and the mean accuracy for the control group was 68% (range: 58% -78%). No significant difference in the mean classification accuracy of pre-neurofeedback offline classifier was observed in both two groups (p=0.820) (as shown in Figure 3A). The average confusion matrices are shown in Figure 3B.

**Figure 3.**
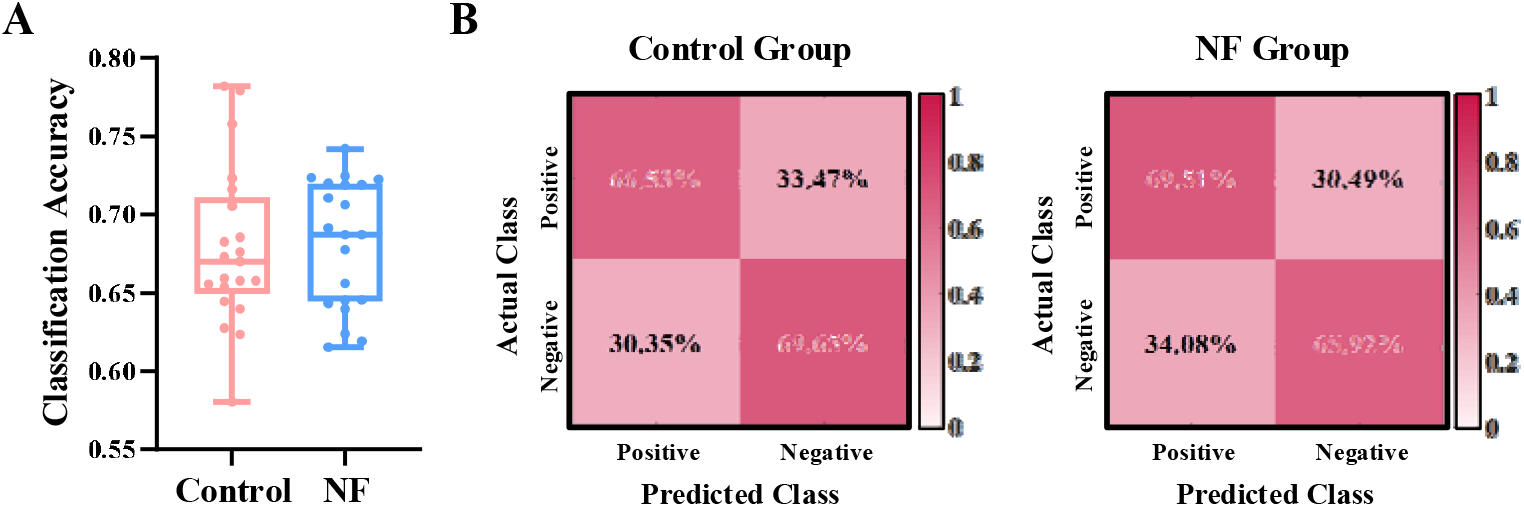
Classification accuracy of the constructed emotion decoder. A. Comparison between the neurofeedback group and control group. B Averaged confusion matrices of each group.

Figure 4 reports the group-level analysis results of the selected informative EEG features for the construction of the emotion classifier. The results show that the top-selected ERP features are the P1 and LPP of occipital-parietal electrodes, N2, P2 and P300 of parietal and central-frontal regions. Among the Hjorth parameter features, the complexity and mobility features of bilateral temporal and anterior frontal electrodes were selected in more subjects. Among PSD features of different frequency bands, beta and gamma-band power features in anterior frontal electrodes were consistently selected. Consistently with PSD features, the more frequently selected DE features are also belong to the high-frequency bands and located in anterior frontal, bilateral temporal, and parietal-occipital regions. Among the wavelet features, the energy of beta and gamma located in the anterior frontal regions was selected with higher frequency.

**Figure 4.**
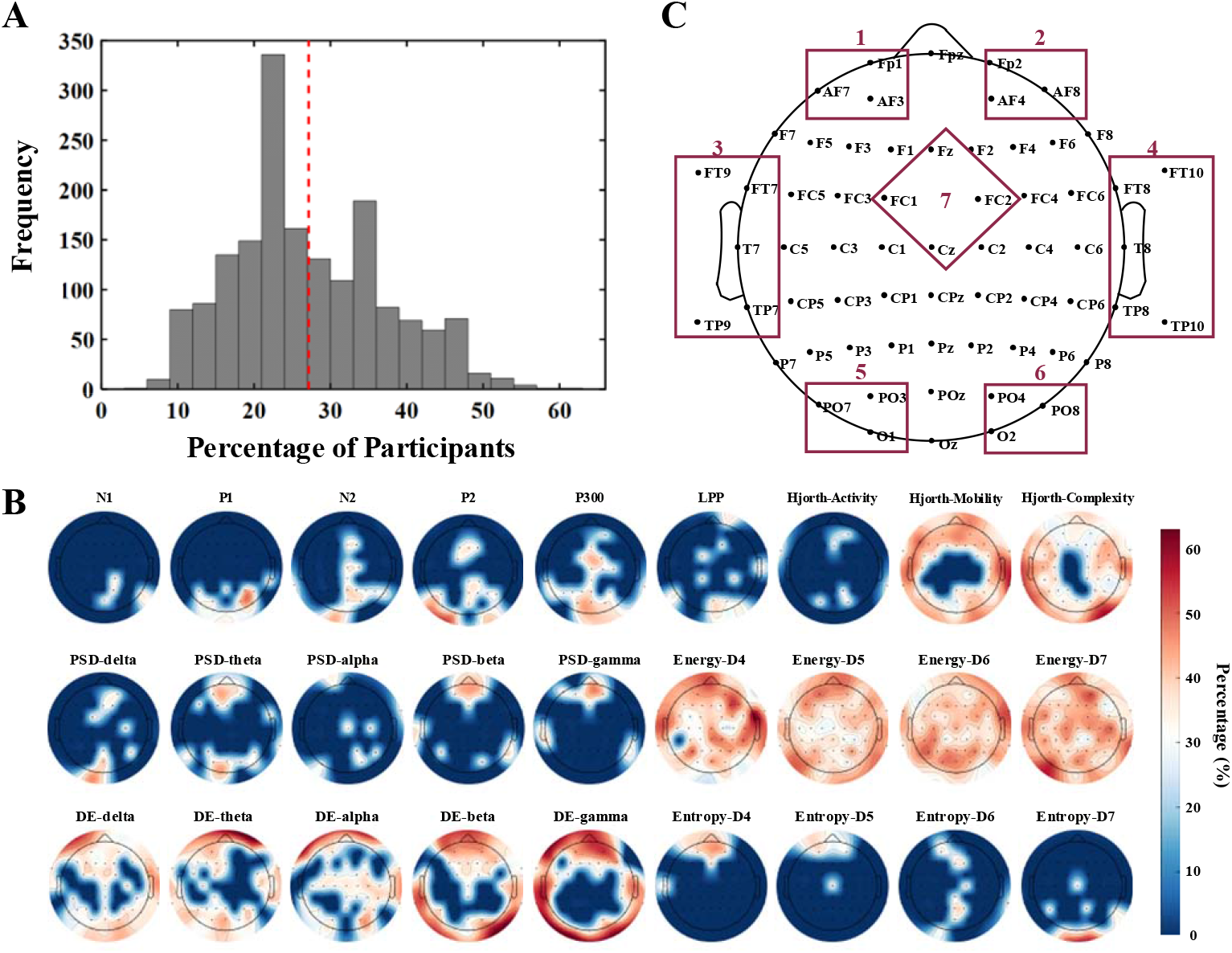
The group-level analysis results of the selected EEG feature for the construction of the emotion classifier. A: The percentage of participants who had each feature selected and used for emotion decoder construction. B: the topographic figures show the features that were selected by more participants (>27%, group mean) at the group level. C. Regions-of-interest (ROIs) for following neural correlates analyses of neurofeedback training.

### 3.4 Neural Correlates of Neurofeedback Training

While neurofeedback training changed the emotional states and showed significant differences in regulation effect between the two groups, we also investigated the alternations of related EEG correlates for the LP regulation run. Therefore, for the selected EEG features (averaged within seven ROIs), paired sample t-tests were applied to make the comparison between the view block and regulation block (Table 2 and Table 3). According to our results, the NF group showed significant changes in ERP potentials of the middle frontal (P300) and parietal regions (P1 and P2), Hjorth mobility of the anterior frontal regions, DE features of the anterior frontal regions (low-frequency rhythm: delta, theta, and alpha), high-frequency Wavelet energy and entropy of the frontal and temporal regions (Beta and Gamma). The Control group showed significant changes in ERP potentials of the middle frontal (P300) and parietal regions (N2 and P2), Hjorth mobility and complexity of the anterior frontal and right temporal regions, DE features of the anterior frontal regions (low-frequency rhythm: delta and theta), high-frequency Wavelet energy of the frontal regions (Beta).

**Table 1.**
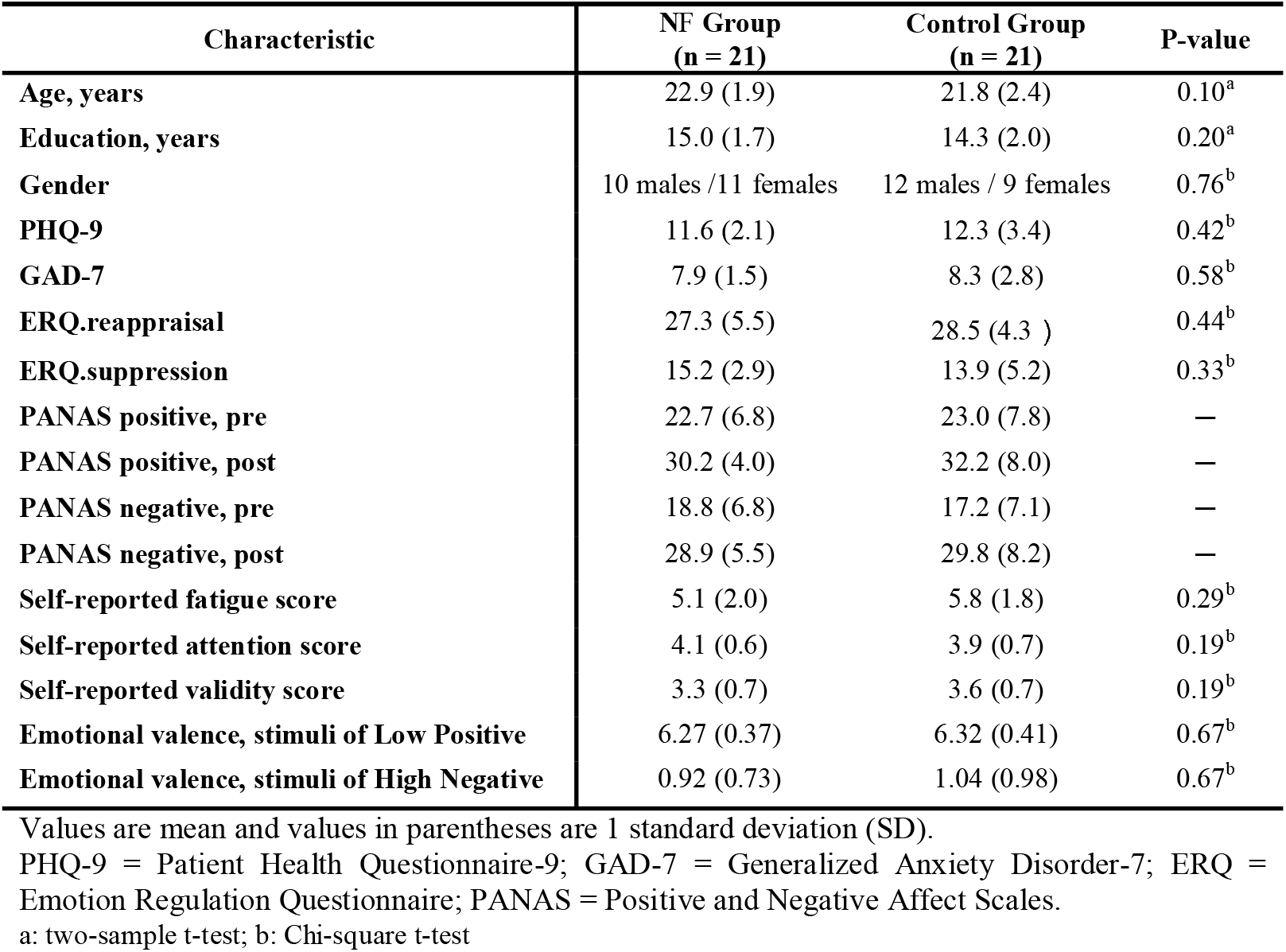
Demographics and behavior ratings.

**Table 2.** Comparison of EEG features between view block and regulation block for the NF group.

| Features | P-value |  |  |  |  |  |  |
| --- | --- | --- | --- | --- | --- | --- | --- |
|  | ROI1 | ROI2 | ROI3 | ROI4 | ROI5 | ROI6 | ROI7 |
| P1 | \ | \ | \ | \ | 0.0005 | \ | \ |
| P2 | \ | \ | \ | \ | \ | 0.0055 | \ |
| P300 | \ | \ | \ | \ | \ | \ | 0.0008 |
| Hjorth-Mobility | 0.0010 | 0.0066 | \ | \ | \ | \ | \ |
| PSD-theta | \ | 0.0051 | \ | \ | \ | \ | \ |
| DE-delta | 0.0004 | 0.0006 | \ | 0.0060 | \ | \ | \ |
| DE-theta | 0.0003 | 0.0002 | \ | \ | \ | \ | \ |
| DE-alpha | \ | 0.0038 | \ | \ | \ | \ | \ |
| DE-beta | \ | 0.0039 | \ | \ | \ | \ | \ |
| DE-gamma | \ | 0.0038 | \ | \ | \ | \ | \ |
| Energy-D4 | 0.0052 | \ | \ | \ | \ | \ | \ |
| Energy-D5 | 0.0009 | 0.0008 | 0.0002 | 0.0056 | \ | \ | \ |
| Entropy-D6 | 0.0046 | \ | \ | \ | \ | \ | \ |

**Table 3.**
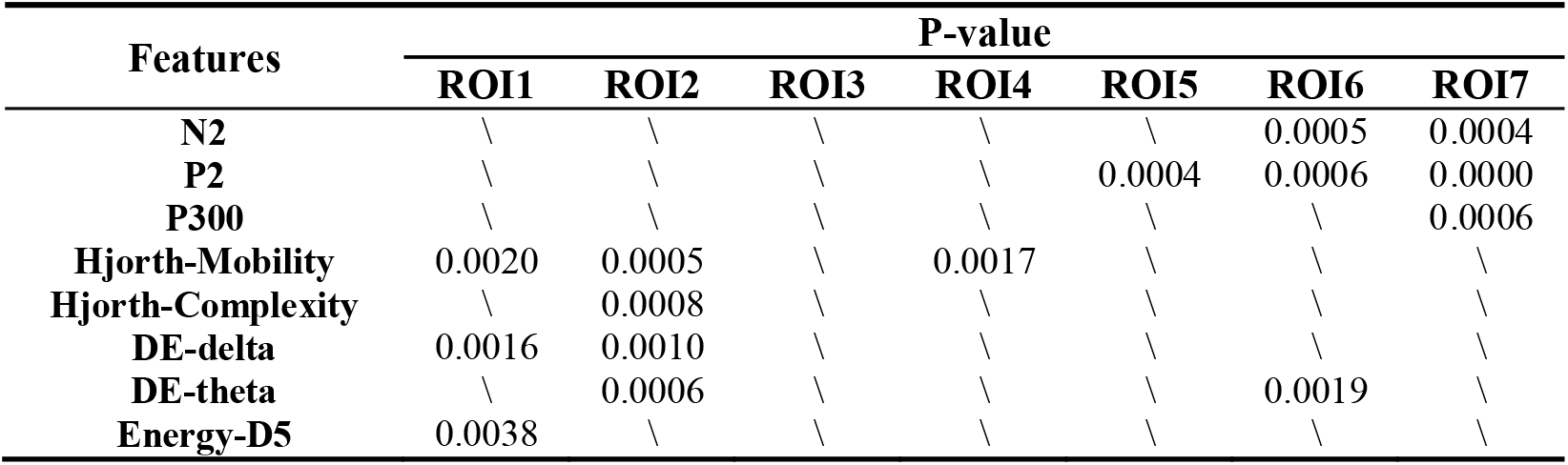
Comparison of EEG features between view block and regulation block for the Control group.

| Features | P-value |  |  |  |  |  |  |
| --- | --- | --- | --- | --- | --- | --- | --- |
|  | ROI1 | ROI2 | ROI3 | ROI4 | ROI5 | ROI6 | ROI7 |
| N2 | \ | \ | \ | \ | \ | 0.0005 | 0.0004 |
| P2 | \ | \ | \ | \ | 0.0004 | 0.0006 | 0.0000 |
| P300 | \ | \ | \ | \ | \ | \ | 0.0006 |
| Hjorth-Mobility | 0.0020 | 0.0005 | \ | 0.0017 | \ | \ | \ |
| Hjorth-Complexity | \ | 0.0008 | \ | \ | \ | \ | \ |
| DE-delta | 0.0016 | 0.0010 | \ | \ | \ | \ | \ |
| DE-theta | \ | 0.0006 | \ | \ | \ | 0.0019 | \ |
| Energy-D5 | 0.0038 | \ | \ | \ | \ | \ | \ |

The results of the Pearson correlation analysis showed that among the selected EEG features in the LP regulation run, significant correlations were observed between regulation-related brain activity changes and neurofeedback training performance (Figure 5). Specifically, the NF group had a significant correlation between the training performance and the amount of change in the high-frequency Wavelet energy of the left temporal region (r=0.616, p=0.003). Besides, the Control group showed significant correlations in the Hjorth mobility of the anterior frontal region (r=0.576, p=0.006).

**Figure 5.**
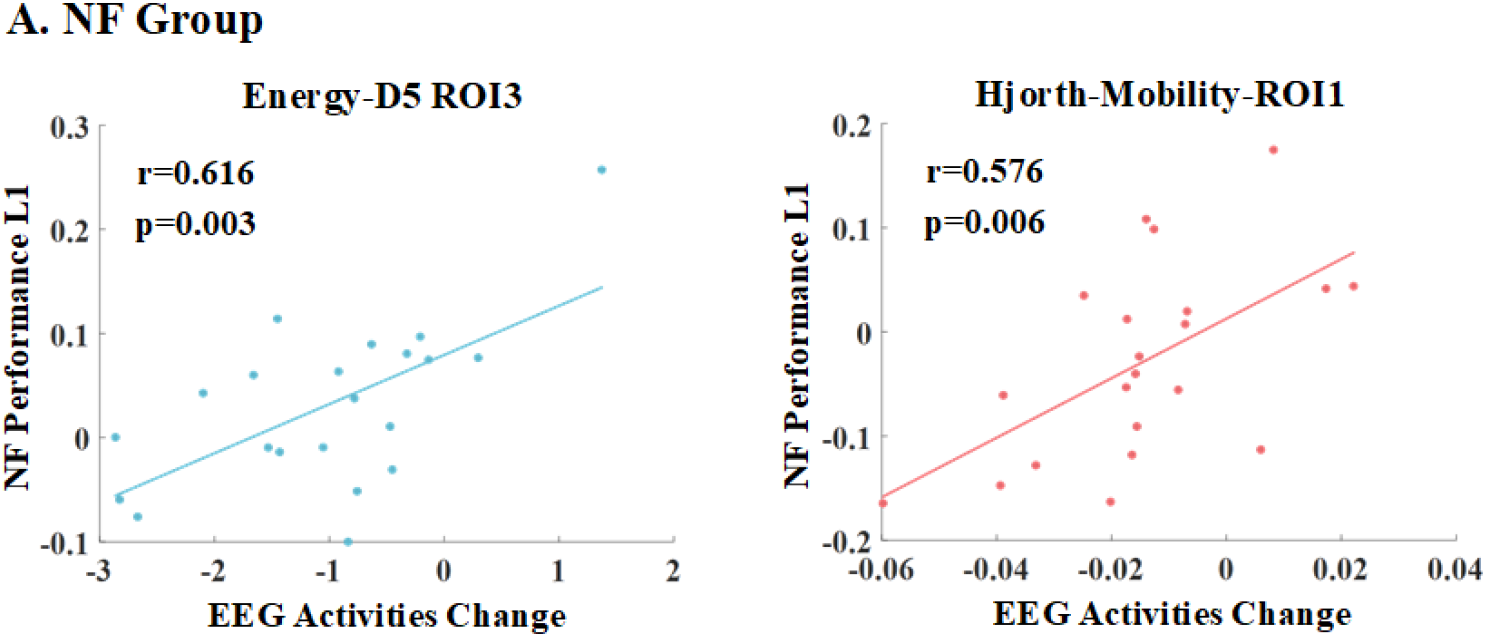
Results of the correlation analysis between the amount of EEG activity changes before and after the neurofeedback and neurofeedback training performance.

## 4 Discussion

### 4.1 Neurofeedback Training Performance for Emotion Regulation

Neurofeedback is an innovative tool and has been used to successfully improve emotion regulation in recent years (Linhartová et al., 2019; Melnikov, 2021). Particularly in light of its relatively low cost, portability, and easy of use compared to fMRI, the use of EEG neurofeedback to improve emotion regulation hold promise for improving results via personalized, targeted protocols. For example, frontal alpha asymmetry EEG neurofeedback has been applied in both healthy (Quaedflieg *et al*., 2016; Li *et al*., 2023) and depressed participants for emotion regulation training (Wang et al., 2019). One recently published study designed a real-time EEG-based brain-computer interface system for neurofeedback training (Huang et al., 2021). The decoded immediate emotional state was provided as a neurofeedback signal to help the subjects update their strategies to regulate their emotion toward a specific emotional state with the help of displayed video clips. A large number of studies have shown that cognitive reappraisal is one of the most flexible, effective, and adaptive regulatory strategies to modulate subjective emotional experience (Webb et al., 2012). A higher reliance on adaptive and effective cognitive reappraisal strategies for emotional regulation is associated with less depression, fewer negative effects, and higher life satisfaction. However, current studies of EEG neurofeedback only target the modulation of spontaneous emotional states and do not involve the cognitive reappraisal of emotionally evocative stimuli. Consequently, it is necessary to develop an EEG neurofeedback paradigm that is more specific to cognitive reappraisal training for emotion regulation.

In this study, we constructed and evaluated a new EEG neurofeedback-guided cognitive reappraisal training protocol. Cognitive reappraisal has two directions of regulation: upregulating positive emotions and downregulating negative emotions, they are both important for managing affective arousal (Gross, 2015). They can be used to cope with adverse situations, and for increasing one’s sense of happiness and have been confirmed to be associated with different neural mechanisms as revealed by fMRI studies (Sokołowski et al., 2022). Previous neurofeedback studies on emotion regulation have only focused on the downregulation of negative emotions (Linhartová et al., 2019; Wang et al., 2019; Zweerings et al., 2020). According to existing literature, cognitive reappraisal has been combined with fMRI neurofeedback training (Zweerings et al., 2020). Thus, research to date does not paint a full picture of whether neurofeedback can help with cognitive reappraisal training. Besides, this study used fMRI signal change between reappraisal regulation and view conditions as the neurofeedback signal for the voluntary control of brain activation in key emotion regulation areas (Zweerings et al., 2020). This kind of neurofeedback signal could not directly reflect the cognitive reappraisal performance. Based on this consideration, we performed real-time emotion classification, calculated the emotion change of the decoded brain state between reappraisal regulation and view conditions, and used it as the neurofeedback signal to inform self-regulation performance. According to our results, the NF group achieved better performance compared with the Control group without feedback for the cognitive reappraisal of emotional stimuli with low positive valence, but not for the cognitive reappraisal of emotional stimuli with high negative valence. This suggested that regulation direction and stimulus valence should be taken into account when applying neurofeedback-enhanced cognitive reappraisal training.

### 4.2 Neural Correlates for Neurofeedback-Guided Cognitive Reappraisal

While emotions are challenging to describe directly, they can be detected in physiological indices, such as EEG. Many researchers working on emotion recognition have used EEG-based methods because signals collected by assessable EEG devices could offer meaning-rich signals with a high temporal resolution. According to the existing literature, emotions can be analyzed using distinguishable features of EEG signals (Liu et al., 2021; Rahman et al., 2021). We analyzed the selected EEG features for emotion classification at the group level, and the wavelet energy features and DE features play a higher important role in emotion classification, especially the frequency components of beta and gamma of the anterior frontal, bilateral temporal, and posterior parietal and occipital regions. Consistent findings were also found in studies concerning the analysis of EEG features in emotion recognition (Wang et al., 2014; Zheng et al., 2019), and it can be explained by that higher frequency brain activities reflect emotional and cognitive processes (Ray and Cole, 1985; Luther et al., 2022). Besides, the relationship between emotion recognition and frontal regions was illustrated in the studies of (Lin et al., 2010). The existing literature also supports the role of the temporal cortex in the processing of visual materials with emotional content (Meletti et al., 2006) and the temporal lobe has been proposed to be informative when it comes to detecting pleasant affect (Kortelainen et al., 2015). For time-domain features, discriminating features of ERP component P300 have been observed over central and posterior regions. Studies have verified the relationship between the amplitude of P300 and the valence of emotional stimuli (Zhu et al., 2015). Hjorth parameter features, the complexity and mobility features, are also the top selected features. This is consistent with the previous studies that claim the importance of Hjorth parameters in EEG classification tasks (Cecchin et al., 2010; Ö et al., 2017).

Neurofeedback is provided to a subject based on his/her brain activity in order to self-regulate his/her brain function. Therefore, changes in the selected features of emotion decoding could reflect the neural plasticity changes induced by emotion regulation. Overall, the two groups of subjects showed relatively consistent changes in EEG features, mainly located in the frontal and parietal regions. For example, decreased P300 amplitude for a more positive perception for both groups, because differences in valence dimensions also can modulate ERP outcomes (Bernat et al., 2001); changes in high-frequency band oscillations were also observed during emotion regulation, because high-frequency-band activities reflect the characteristics of emotion integration, particularly in the cognitive control of emotions (Tang et al., 2011), and pleasant pictures elicit stronger responses than other stimuli in the high gamma band (Boucher et al., 2015). Significant correlations between NF learning performance and changes in EEG activities were observed in the high-frequency Wavelet energy of the left temporal region for the NF group, and the Hjorth mobility of the anterior frontal region for the control group. Cognitive reappraisal is the process of reinterpreting an emotional stimulus to change the emotional response to that stimulus in order to achieve emotion regulation, so language and memory processes are also key to the successful implementation of cognitive reappraisal. The frontal lobes and emotions are closely linked and are key brain regions for emotion regulation, where activity in the left frontal lobe is associated with approach motivation and positive emotions, while activity in the right frontal lobe is associated with avoidance motivation and negative emotions (Baehr et al., 1997). The temporal lobe constitutes the semantic system (Binder et al., 2009) and can help store and retrieve relevant information (Jackson, 2021), and many studies have repeatedly found activation of temporal lobe areas of the brain in cognitive reappraisal (Goldin et al., 2008; Kanske et al., 2011; Dörfel et al., 2014). The differences exhibited here by the two groups may lie in different cognitive reappraisal strategies adopted during neurofeedback training.

## Limitations and Future Directions

The present study has several limitations and the findings still need further validation. First, the single-session neurofeedback training design was applied and the training performance was assessed within a short period. Prior research in this area has focused on training performance across multiple training sessions for both self-regulation of EEG activities and the long-term effects of enhanced behavioral performance (Bu et al., 2019; Huang et al., 2021). Second, considering gender differences in emotion regulation (Nolen-Hoeksema and Aldao, 2011), future studies with larger sample sizes could report data analysis results separately for male and female participants. Third, it is unclear how this decoded EEG NF-guided cognitive reappraisal training paradigm can be transferred to patient groups, such as patients with major depression. A potential line of research is to further verify the ability of this training paradigm in patients with emotion-related mental disorders.

## Conclusions

In conclusion, we developed and tested a novel decoded EEG neurofeedback-guided cognitive reappraisal emotion training protocol. Real-time feedback of the regulation effect helps subjects improve emotion regulation performance for emotional stimuli with low positive valence. This neurofeedback training protocol is a promising treatment for emotion-related mental disorders, with the potential to be a low-cost and high-portability brain-based, non-invasive, neural modulation technique.

## Acknowledgments

We would like to extend our thanks to the adolescents who participated in this study and their parents. This work was supported by the National Natural Science Foundation of China (No. 62276169 and 82272114), Shenzhen Soft Science Research Program Project (No. RKX20220705152815035), Shenzhen Science and Technology Research and Development Fund for Sustainable Development Project (No. KCXFZ20201221173613036).

## Ethics Approval

The study protocol was approved by the Medical Ethics Committee of Shenzhen University. Informed, written consent was obtained from all participants and their caregivers prior to participation in the study.

